## Supplementary Figures and Tables S1 to S4 for "The presence of copy number variants in specific topologically associating domains has prognostic value in many cancer types"

This document contains

**Supplementary Figures S1 to S8**

**Supplementary Tables S1 to S4**

### Supplementary Figures

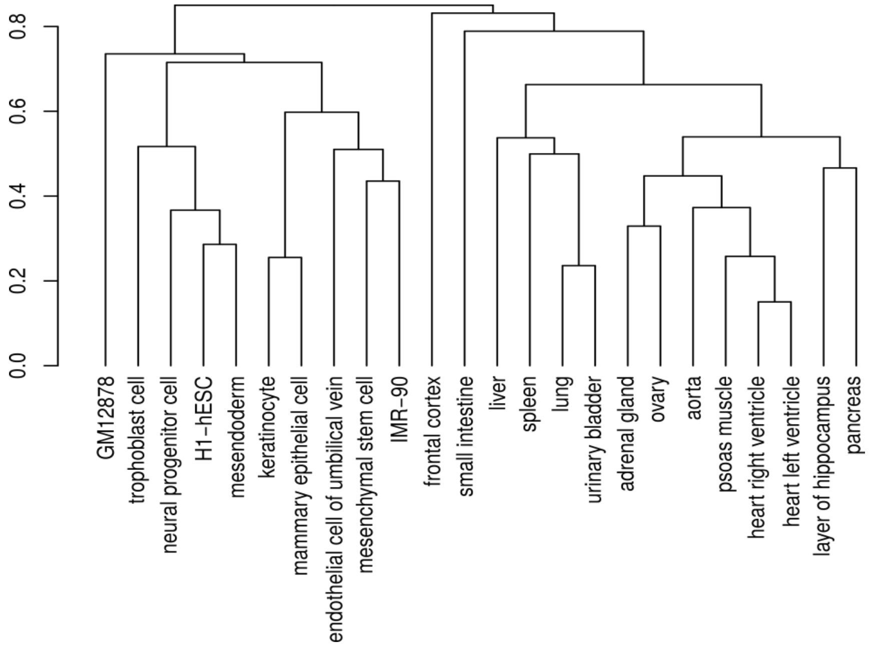


**Figure S1. Relationship among 24 normal tissues, based on the similarity of their expression profiles.**

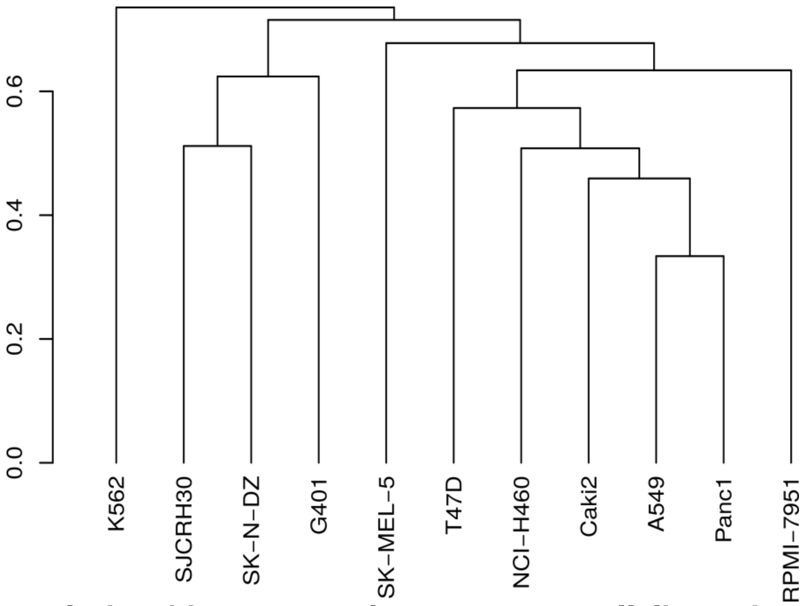


**Figure S2. Relationship among eleven cancer cell lines, based on the similarity of their expression profiles.**

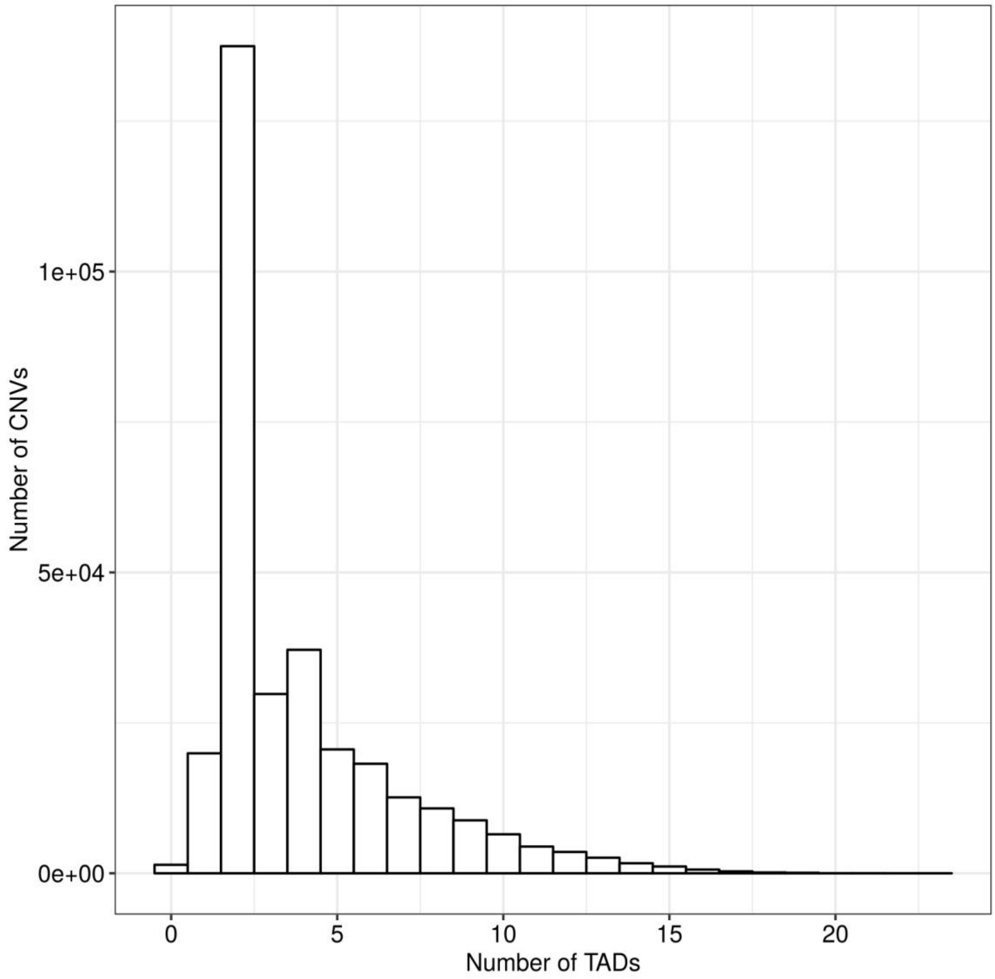


**Figure S3. Number of TADs overlapping with each CNV.**

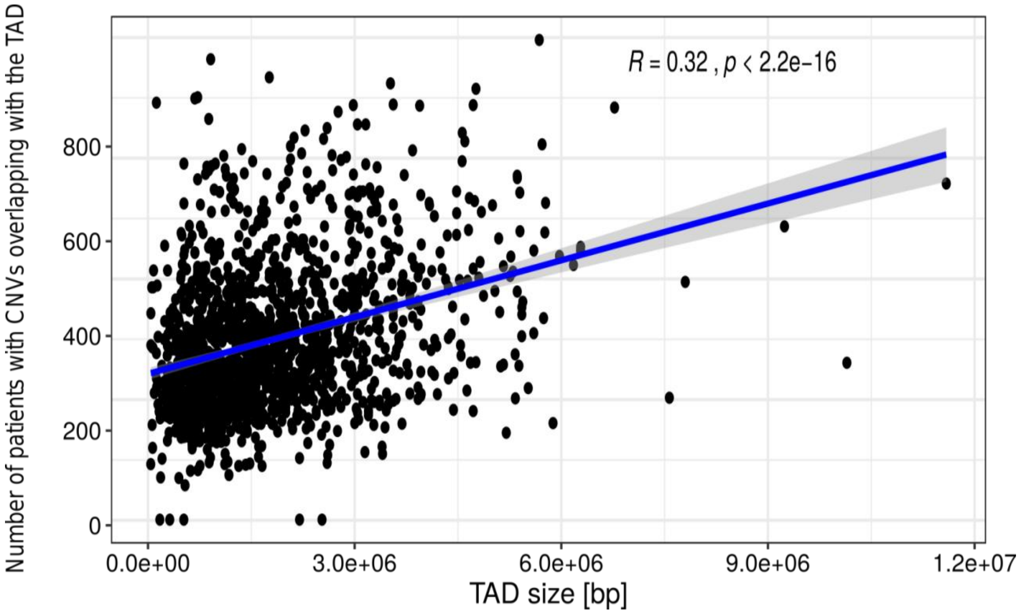


**Figure S4. The size of the TADs and the number of CNVs overlapping with the TADs are only weakly correlated.** Number of patients with one or more CNVs overlapping with a TAD as a function of the size of the TAD. The blue line is fitted linear model describing the relationship between TAD size and number of CNVs overlapping with the TAD. The grey shade represents the 95% confidence interval of the fitted value. R is the Spearman’s correlation coefficient and p is the P-value for Spearman’s correlation test.

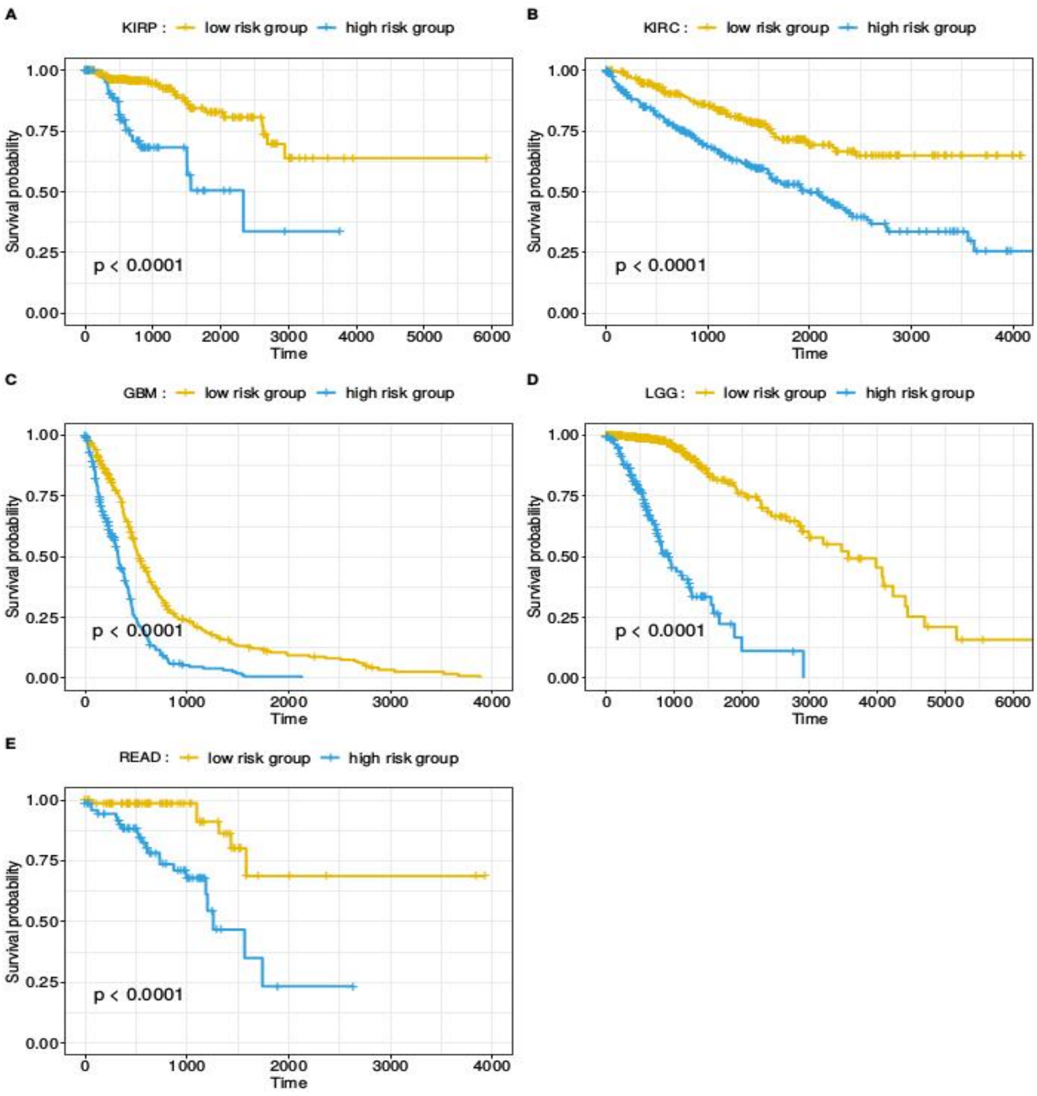


**Figure S5. Kaplan-Meier analysis for KIRP, KIRC, GBM, LGG AND READ.** Patients were separated into high- (blue) and a low-risk (yellow) groups according to the prognostic features of the final LASSO Cox regression model and subjected to Kaplan-Meier analysis. A) KIRP; B) KIRC; C) GBM; D) LGG; E) READ.

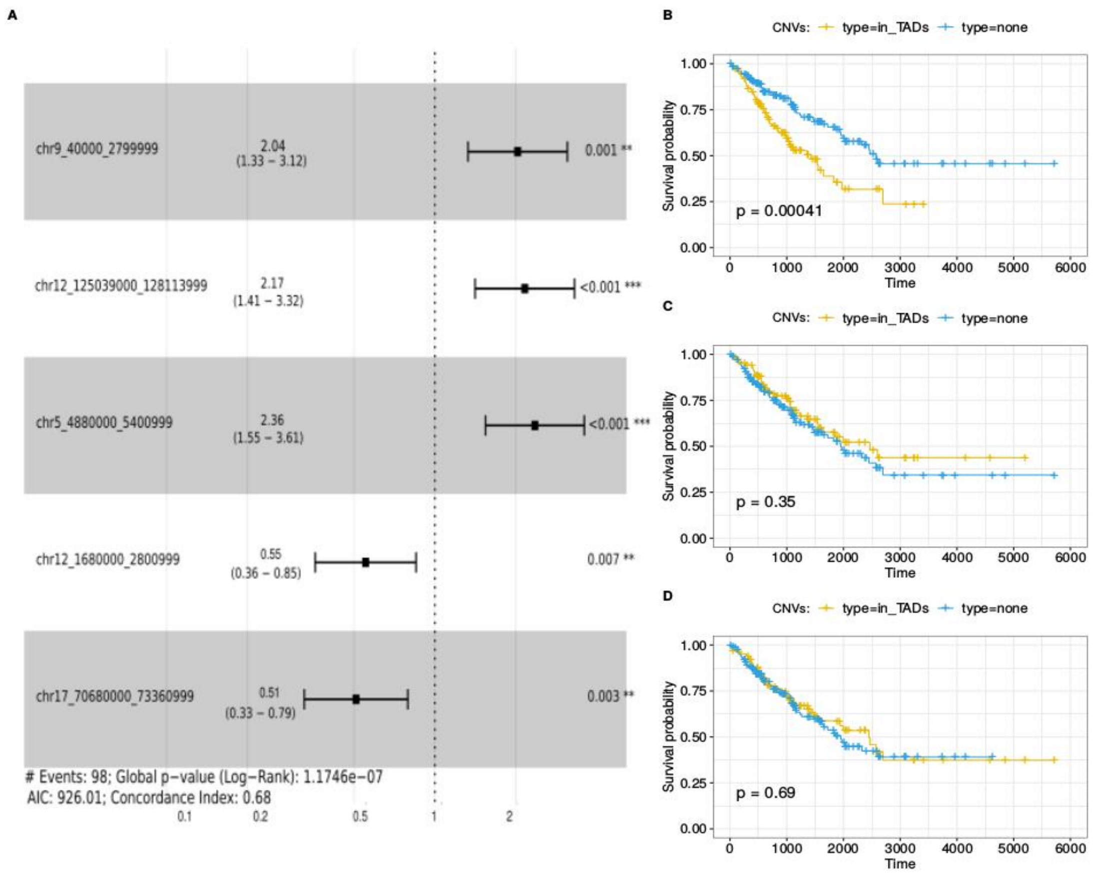


**Figure S6. Survival analysis for SARC patients. A)** Hazard ratios with 95% confidence intervals for all prognostic (P-values < 0.05) TADs from the final LASSO Cox regression model. **B-D)** Kaplan-Meier curves for patients separated according to the presence or absence of CNVs in prognostic TADs: **B)** chr5:4880000-5400999; **C)** chr12:1680000-2800999; and **D)** chr17:70680000-73360999.

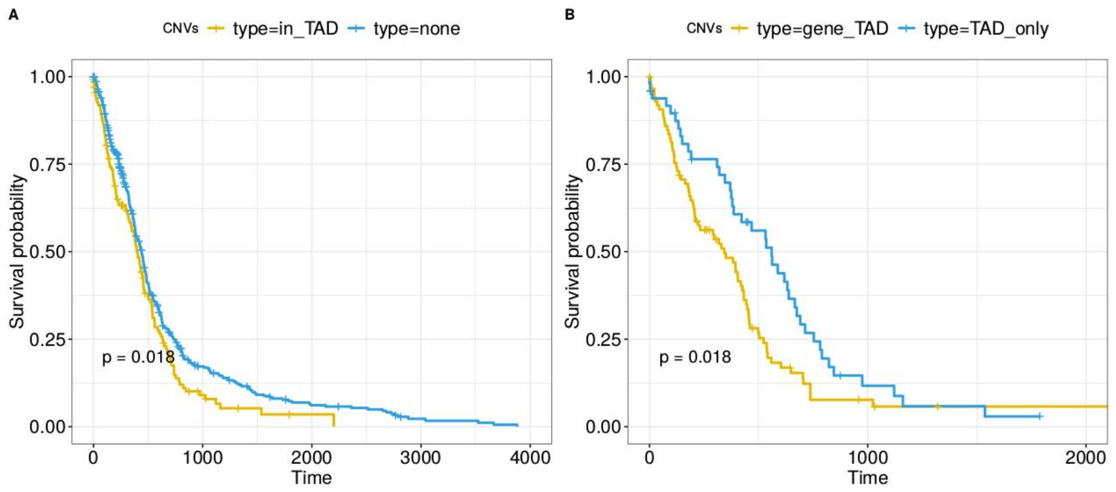


**Figure S7. Kaplan-Meier survival analysis for GBM patients separated according to the presence or absence of CNVs in the prognostic TAD chr12:55679000-57720999. A)** Survival of the 136 patients that had CNVs in the TAD (yellow) compared to that of the 453 patients that did not (blue). **B)** Survival of the 87 patients that had CNVs in *DDIT3* (yellow) compared to that of the 49 patients that had CNVs in the TAD but not in *DDIT3* (blue).

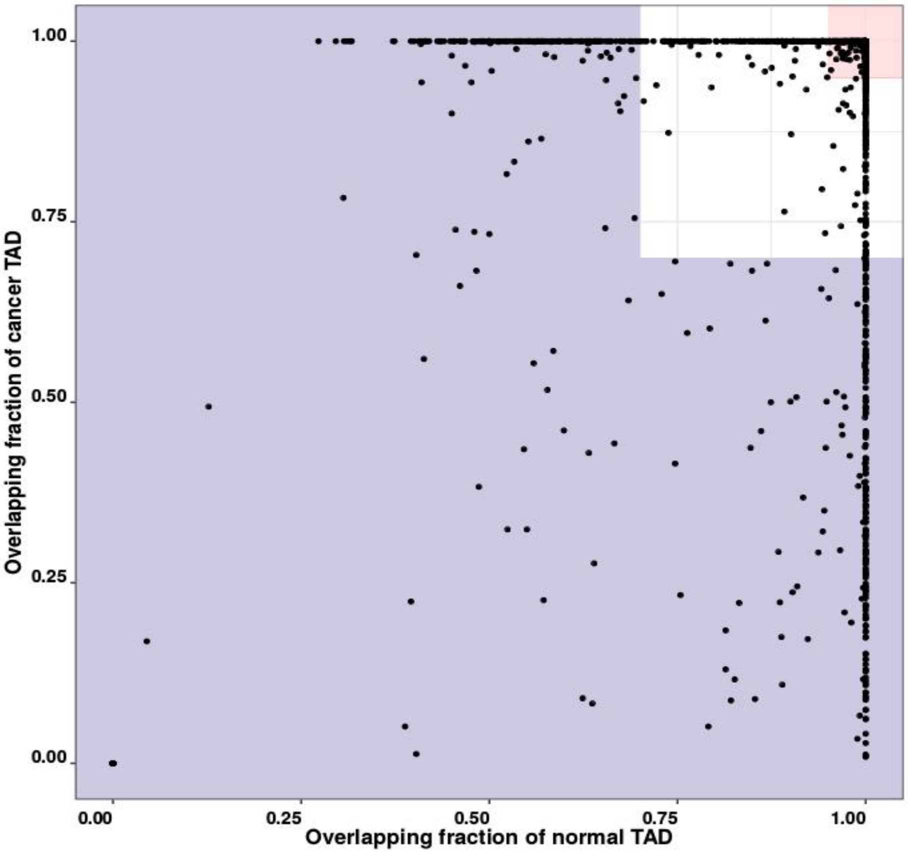


**Figure S8. Fraction of mutual overlaps for normal and cancer TADs.** Each dot represents a normal TAD. The x-axis shows the fraction of the sequence of the (normal) TAD that overlapped with a cancer TAD; if a (normal) TAD overlapped with multiple cancer TADs, we selected the TAD with the highest fraction. For the corresponding normal TAD, the y-axis shows the fraction of its sequence overlapping with the normal TAD. Constitutive TADs are highlighted in red; non-constitutive TADs are shown in purple; the remaining TADs were considered "ambiguous".

### Supplementary Tables

**Supplementary Table S1. Number of TADs in each TAD map.** TAD maps were retrieved for 25 normal tissues/cell lines and 12 cancer cell lines.

| Tissue/Cell line | State | Description | Number of TADs |
|---|---|---|---|
| **Adrenal** | Normal | Adrenal gland | 1,408 |
| **Aorta** | Normal | Aorta | 1,564 |
| **Bladder** | Normal | Urinary bladder | 1,416 |
| **Bowel Small** | Normal | Small intestine | 1,276 |
| **Cortex DLPFC** | Normal | Frontal cortex | 1,469 |
| **GM12878_Lieberman** | Normal | *Lymphoblastoid* cell line | 1,309 |
| **GM12878_Rao** | Normal | *Lymphoblastoid* cell line | 3,255 |
| **H1-ESC** | Normal | H1 human embryonic stem cell line | 2,414 |
| **H1-MES** | Normal | Mesendoderm | 2,442 |
| **H1-MSC** | Normal | Mesenchymal stem cell | 2,295 |
| **H1-NPC** | Normal | Neural progenitor cell | 1,676 |
| **H1-TRO** | Normal | Trophoblast cell | 2,056 |
| **Hippocampus** | Normal | Layer of hippocampus | 1,504 |
| **HMEC** | Normal | Mammary epithelial cell | 3,250 |
| **HUVEC** | Normal | Endothelial cell of umbilical vein | 2,468 |
| **IMR90** | Normal | Fetal lung cells | 3,094 |
| **Liver** | Normal | Liver | 1,827 |
| **Lung** | Normal | Lung | 1,438 |
| **NHEK** | Normal | Keratinocyte | 3,177 |
| **Ovary** | Normal | Ovary | 1,222 |
| **Pancreas** | Normal | Pancreas | 1,397 |
| **Psoas** | Normal | Psoas muscle | 1,335 |
| **Spleen** | Normal | Spleen | 2,432 |
| **Ventricle Left** | Normal | Heart left ventricle | 1,738 |
| **Ventricle Right** | Normal | Heart right ventricle | 1,635 |
| **A549** | Cancer | Lung adenocarcinoma | 1,866 |
| **Caki2** | Cancer | Papillary renal cell carcinoma | 1,919 |
| **K562_Lieberman** | Cancer | Chronic myelogenous leukemia | 1,480 |
| **K562_Rao** | Cancer | Chronic myelogenous leukemia | 3,243 |
| **NCIH460** | Cancer | Large cell lung carcinoma | 1,664 |
| **PANC1** | Cancer | Pancreatic ductal adenocarcinoma | 1,847 |
| **RPMI7951** | Cancer | Melanoma | 1,764 |
| **SJCRH30** | Cancer | Metastatic rhabdomyosarcoma | 1,387 |
| **T470** | Cancer | | 1,889 |
| **G401** | Cancer | Rhabdoid tumor of the kidney | 1,984 |
| **SKNDZ** | Cancer | Invasive ductal carcinoma | 1,515 |

| | | | |
|---|---|---|---|
| **SKMEL5** | Cancer | Cutaneous melanoma | 2,256 |

**Supplementary Table S2. RNA-sequencing data used for the computation of the weights summarizing the contribution of the individual TAD maps to the consensus TADs.**

| Tissues/cell lines | Assay |
|---|---|
| **A549** | Poly-A RNA-seq |
| **aorta** | Poly-A RNA-seq |
| **endothelial cell of umbilical vein** | Poly-A RNA-seq |
| **H1-hESC** | Poly-A RNA-seq |
| **heart right ventricle** | Poly-A RNA-seq |
| **keratinocyte** | Poly-A RNA-seq |
| **layer of hippocampus** | Poly-A RNA-seq |
| **mesenchymal stem cell** | Poly-A RNA-seq |
| **mesendoderm** | Poly-A RNA-seq |
| **neural stem progenitor cell** | Poly-A RNA-seq |
| **pancreas** | Poly-A RNA-seq |
| **psoas muscle** | Poly-A RNA-seq |
| **small intestine** | Poly-A RNA-seq |
| **trophoblast cell** | Poly-A RNA-seq |
| **adrenal gland** | Total RNA-seq |
| **Caki2** | Total RNA-seq |
| **frontal cortex** | Total RNA-seq |
| **G401** | Total RNA-seq |
| **GM12878** | Total RNA-seq |
| **heart left ventricle** | Total RNA-seq |
| **IMR-90** | Total RNA-seq |
| **K562** | Total RNA-seq |
| **liver** | Total RNA-seq |
| **lung** | Total RNA-seq |
| **mammary epithelial cell** | Total RNA-seq |
| **mesenchymal stem cell of the bone marrow** | Total RNA-seq |
| **NCI-H460** | Total RNA-seq |
| **neural progenitor cell** | Total RNA-seq |
| **ovary** | Total RNA-seq |
| **RPMI-7951** | Total RNA-seq |
| **SJCRH30** | Total RNA-seq |
| **SK-MEL-5** | Total RNA-seq |
| **SK-N-DZ** | Total RNA-seq |
| **spleen** | Total RNA-seq |
| **urinary bladder** | Total RNA-seq |
| **T47D** | Poly-A RNA-seq |
| **Panc1** | Poly-A RNA-seq |

**Supplementary Table S3. Summary of TCGA patients included in the analysis.** Third column indicated the total number of patients with CNVs from TCGA for each of 32 cancer types; fourth column referred to the number of patients after filtering (see methods) and fifth column indicated the cancer types applied in survival analysis or not.

| Project simple id | Disease | Number of patients (pre-filtering) | Number of patients (post-filtering) | Survival Analysis |
|---|---|---|---|---|
| **ACC** | Adrenocortical carcinoma | 90 | 90 | No |
| **BLCA** | Bladder Urothelial Carcinoma | 412 | 412 | Yes |
| **BRCA** | Breast invasive carcinoma | 1094 | 1088 | Yes |
| **CESC** | Cervical squamous cell carcinoma and endocervical adenocarcinoma | 295 | 295 | Yes |
| **CHOL** | Cholangiocarcinoma | 36 | 36 | No |
| **COAD** | Colon adenocarcinoma | 450 | 449 | Yes |
| **DLBC** | Lymphoid Neoplasm Diffuse Large B-cell Lymphoma | 48 | 48 | No |
| **ESCA** | Esophageal carcinoma | 184 | 184 | Yes |
| **GBM** | Glioblastoma multiforme | 590 | 589 | Yes |
| **HNSC** | Head and Neck squamous cell carcinoma | 517 | 517 | Yes |
| **KICH** | Kidney Chromophobe | 66 | 66 | |
| **KIRC** | Kidney renal clear cell carcinoma | 530 | 522 | Yes |
| **KIRP** | Kidney renal papillary cell carcinoma | 290 | 289 | Yes |
| **LGG** | Brain Lower Grade Glioma | 514 | 510 | Yes |
| **LIHC** | Liver hepatocellular carcinoma | 375 | 374 | Yes |
| **LUAD** | Lung adenocarcinoma | 518 | 516 | Yes |
| **LUSC** | Lung squamous cell carcinoma | 503 | 502 | Yes |
| **MESO** | Mesothelioma | 87 | 87 | No |
| **OV** | Ovarian serous cystadenocarcinoma | 568 | 568 | Yes |
| **PAAD** | Pancreatic adenocarcinoma | 184 | 184 | Yes |
| **PCPG** | Pheochromocytoma and Paraganglioma | 178 | 175 | No |
| **PRAD** | Prostate adenocarcinoma | 497 | 493 | No |
| **READ** | Rectum adenocarcinoma | 164 | 164 | Yes |
| **SARC** | Sarcoma | 260 | 258 | Yes |
| **SKCM** | Skin Cutaneous Melanoma | 104 | 103 | no |
| **STAD** | Stomach adenocarcinoma | 442 | 440 | Yes |
| **TGCT** | Testicular Germ Cell Tumors | 134 | 134 | No |
| **THCA** | Thyroid carcinoma | 505 | 446 | No |
| **THYM** | Thymoma | 124 | 118 | No |
| **UCEC** | Uterine Corpus Endometrial Carcinoma | 540 | 529 | Yes |
| **UCS** | Uterine Carcinosarcoma | 56 | 56 | No |
| **UVM** | Uveal Melanoma | 80 | 79 | No |

**Supplementary Table 4. Functional analysis for 79 TADs enriched with CNVs.**

| Category | Term | Count | P-value | List total | Pop Hits | Pop total | **FDR** |
|---|---|---|---|---|---|---|---|
| **GOTERM_MF_DIRECT** | GO:0004519~endonuclease activity | 19 | $7.5x10^{-08}$ | 1436 | 44 | 14331 | $1.2x10^{-06}$ |
| **GOTERM_MF_DIRECT** | GO:0004540~ribonuclease activity | 12 | $7.0x10^{-07}$ | 1436 | 20 | 14331 | $1.1x10^{-05}$ |
| **GOTERM_MF_DIRECT** | GO:0005132~type I interferon receptor binding | 7 | $2.3x10^{-05}$ | 1436 | 8 | 14331 | $3.8x10^{-04}$ |
| **GOTERM_CC_DIRECT** | GO:0045095~keratin filament | 24 | $4.5x10^{-05}$ | 1555 | 94 | 15378 | $6.8x10^{-04}$ |
| **GOTERM_CC_DIRECT** | GO:0005796~Golgi lumen | 22 | $6.8x10^{-05}$ | 1555 | 84 | 15378 | $1.0x10^{-03}$ |
| **KEGG_PATHWAY** | hsa04623:Cytosolic DNA-sensing pathway | 15 | $1.2x10^{-04}$ | 566 | 48 | 5840 | $1.5x10^{-03}$ |
| **GOTERM_BP_DIRECT** | GO:0002323~natural killer cell activation involved in immune response | 7 | $1.5x10^{-04}$ | 1426 | 10 | 14258 | $2.7x10^{-03}$ |
| **GOTERM_BP_DIRECT** | GO:0033141~positive regulation of peptidyl-serine phosphorylation of STAT protein | 7 | $1.5x10^{-04}$ | 1426 | 10 | 14258 | $2.7x10^{-03}$ |
| **GOTERM_BP_DIRECT** | GO:0060337~type I interferon signaling pathway | 16 | $1.5x10^{-04}$ | 1426 | 53 | 14258 | $2.7x10^{-03}$ |
| **GOTERM_BP_DIRECT** | GO:0002286~T cell activation involved in immune response | 7 | $5.4x10^{-04}$ | 1426 | 12 | 14258 | $9.9x10^{-03}$ |
| **GOTERM_BP_DIRECT** | GO:0050907~detection of chemical stimulus involved in sensory perception | 17 | $1.0x10^{-03}$ | 1426 | 69 | 14258 | $1.9x10^{-02}$ |
| **KEGG_PATHWAY** | hsa05160:Hepatitis C | 23 | $1.6x10^{-03}$ | 566 | 117 | 5840 | $2.1x10^{-02}$ |
| **GOTERM_BP_DIRECT** | GO:0045087~innate immune response | 52 | $1.7x10^{-03}$ | 1426 | 337 | 14258 | $3.1x10^{-02}$ |
| **GOTERM_BP_DIRECT** | GO:0090502~RNA phosphodiester bond hydrolysis, endonucleolytic | 11 | $2.2x10^{-03}$ | 1426 | 36 | 14258 | $3.9x10^{-02}$ |
